# Differential Effects of Extended Exercise and Memantine Treatment on Adult Neurogenesis in Male and Female Rats

**DOI:** 10.1101/332890

**Authors:** Shaina P Cahill, John Darby Cole, Ru Qi Yu, Jack Clemans-Gibbon, Jason S Snyder

## Abstract

The creation of new neurons in adulthood has potential for treating a number of disorders that are characterized by neurodegeneration or impaired plasticity. Animal models of reduced neurogenesis, and studies of the volume and structural integrity of the hippocampus in humans, suggest a possible therapeutic role for adult neurogenesis in age-related cognitive decline, depression, and schizophrenia. Research over the past 20 years has identified a number of approaches for enhancing adult neurogenesis, such as exercise, NMDA receptor antagonists, antidepressant drugs and environmental enrichment. However, despite the chronic nature of many disorders that impact the human hippocampus, most animal studies have only examined the efficacy of neurogenic treatments over relatively short timescales (∼1 month or less). Additionally, investigations into the regulation of neurogenesis typically include only 1 sex, even though many disorders that affect the hippocampus differentially impact males and females. Here, we therefore tested whether two known pro-neurogenic treatments, running and the NMDA receptor antagonist, memantine, could lead to long-term increases in neurogenesis in male and female rats. We found that continuous access to a running wheel (cRUN) initially increased neurogenesis in both sexes, but effects were minimal after 1 month (both sexes) and completely absent after 5 months (males). Similarly, a single injection of memantine (sMEM) only transiently increased adult neurogenesis in both males and females. To determine whether extended increases in neurogenesis were possible with 2 months of RUN and MEM treatments, we subjected rats to interval running (iRUN), weekly memantine injections (mMEM), or combined treatments (iRUN-mMEM, mMEM-iRUN). We found that 2 months of iRUN increased DCX^+^ cell density in females but iRUN-mMEM treatment increased DCX^+^ cell density in males. However, analyses with thymidine analogs revealed that neurogenesis was minimally increased during the initial phases of the 2 month treatments. Collectively, our findings identify sex differences in the efficacy of neurogenic manipulations, which may be relevant for designing plasticity-promoting treatments that target the hippocampus.

## INTRODUCTION

In the dentate gyrus subregion of the hippocampus, adult-born neurons have enhanced plasticity and unique connectivity relative to older neurons (Snyder & Cameron, 2012; Toni & Schinder, 2015), and they play an important role in memory and emotional behavior (Abrous & Wojtowicz, 2015; Cameron & Glover, 2015). Their functional role has stimulated much research on regulatory factors that could be harnessed, typically to promote neurogenesis, in order to enhance cognitive function or recovery from neurological disorders (Toda, Parylak, Linker, & Gage, 2018). However, most studies have examined neurogenesis regulation over hours, days or weeks. Since most disorders of hippocampal function are chronic, it is important to identify whether neurogenesis can be increased over extended intervals to potentially offset long-term dysfunction (e.g. months in rodents, years in humans).

In humans, reduced hippocampal volume is typically interpreted as a sign of damage and is observed in a number of disorders including depression (McKinnon, Yucel, Nazarov, & MacQueen, 2009), schizophrenia (Harrison, 2004), mild cognitive impairment (Yassa et al., 2010), and Alzheimer’s disease (Jack et al., 2000). While the mechanisms underlying hippocampal volume changes are multifaceted, and certainly not wholly reflective of neurogenesis (Schoenfeld, McCausland, Morris, Padmanaban, & Cameron, 2017), changes in adult neurogenesis could contribute to structural damage as well as recovery. Neurogenesis is difficult to measure in humans, and currently can only be assessed in post mortem tissue. However, a number of known neurogenic treatments (identified in animal studies) are associated with reversal of hippocampal structural deficits in humans. Antidepressant treatment restores dentate gyrus granule cell number in depressed patients (Boldrini et al., 2013; Mahar et al., 2017), possibly by increasing adult neurogenesis (Boldrini et al., 2009). Exercise increases hippocampal volume in healthy individuals (Erickson et al., 2011; Killgore, Olson, & Weber, 2013), women with mild cognitive impairment (Brinke et al., 2015) and schizophrenic patients (Pajonk et al., 2010), though effects in schizophrenia have been inconsistent (Kim, Barr, Honer, & Procyshyn, 2018). Moreover, there are sex differences in the prevalence of disorders that impact the hippocampus, with depression (Bangasser & Valentino, 2014; Seedat et al., 2009) and Alzheimer’s disease (Gao, Hendrie, Hall, & Hui, 1998) more common in females, and schizophrenia more common in males (Aleman, Kahn, & Selten, 2003). Thus, it is important to determine which factors can effectively increase adult neurogenesis in males and females, and potentially improve behavioral outcomes.

Neurogenesis is a multistep process whereby precursor cells undergo lineage-directed cell division to produce immature neurons, of which only a fraction is selected to survive and contribute to hippocampal function. Animal models have identified a number of factors that increase neurogenesis, either by promoting precursor proliferation or enhancing immature neuronal survival. For example, factors that can increase proliferation include exercise (Eadie, Redila, & Christie, 2005; Kronenberg et al., 2006), antidepressant drugs and electroconvulsive shock (Malberg, Eisch, Nestler, & Duman, 2000), synthetic chemicals (Petrik et al., 2012), learning (Dupret et al., 2007) and NMDA receptor antagonists (Cameron, McEwen, & Gould, 1995; Maekawa et al., 2009). Survival of immature neurons is enhanced by exercise (Snyder, Glover, Sanzone, Kamhi, & Cameron, 2009) and learning (Dupret et al., 2007; Epp, Spritzer, & Galea, 2007; Gould, Beylin, Tanapat, Reeves, & Shors, 1999). Importantly, regulatory factors can be highly dose and time-dependent, with some factors increasing or decreasing neurogenesis depending on the conditions (e.g. similar experiences can increase or decrease neurogenesis depending on the age of the cell and the extent of learning (Dupret et al., 2007; Olariu, Cleaver, Shore, Brewer, & Cameron, 2005).)

Here we focus on two treatments that have been shown to increase neurogenesis in rodents: running (RUN) and memantine (MEM). RUN is likely the most well-studied method for increasing neurogenesis. In addition to increasing proliferation and survival, it also increases the dendritic complexity of newborn neurons, promotes spine formation, and accelerates their functional maturation (Dostes et al., 2016; Piatti et al., 2011; Redila & Christie, 2006; van Praag, Kempermann, & Gage, 1999; Vivar, Peterson, & van Praag, 2015). However, RUN does not increase neurogenesis in socially isolated rats (Leasure & Decker, 2009; Stranahan, Khalil, & Gould, 2006), in wild mice (Hauser, Klaus, Lipp, & Amrein, 2009), in mice that run only in the light phase (Holmes, Galea, Mistlberger, & Kempermann, 2004), or in animals that run at high intensities (Grégoire, Bonenfant, Le Nguyen, Aumont, & Fernandes, 2014; Naylor, Persson, Eriksson, Jonsdottir, & Thorlin, 2005; So et al., 2017). Furthermore, a number of studies have found that RUN-induced increases in neurogenesis are transient, raising questions about the extent to which it may be used as a strategy for long-term enhanced production of new neurons (Clark et al., 2010; Kronenberg et al., 2006; Naylor et al., 2005; Snyder et al., 2009). Interestingly, there is evidence that neurogenesis may be sustained for extended durations if RUN amount is restricted (Naylor et al., 2005; Nguemeni et al., 2018). However, since these studies only investigated cells born at a single timepoint in male rats, it remains unclear whether restricted RUN enhances neurogenesis consistently across sexes and over extended periods of time.

MEM is a low-affinity NMDA receptor antagonist that has neuroprotective effects and has been approved as an Alzheimer’s disease treatment (Lipton, 2004). In mice, MEM increases cellular proliferation, the size of the stem cell pool, and the production of new neurons by 2-3 fold (Akers et al., 2014; Ishikawa et al., 2014; Maekawa et al., 2009; Namba, Maekawa, Yuasa, Kohsaka, & Uchino, 2009). Notably, two other NMDA receptor antagonists, MK-801 and ketamine, have also been found to increase cell proliferation in the dentate gyrus (Cameron et al., 1995; Soumier, Carter, Schoenfeld, & Cameron, 2016). While repeat dosing of MEM has been found to broadly increase neurogenesis (Ishikawa, Fukushima, Frankland, & Kida, 2016), the efficacy of single vs. multiple doses of MEM remain unknown.

Here, we investigated the long-term efficacy of neurogenic treatments in male and female rats. Rats were subjected to RUN, MEM, or alternating blocks of RUN and MEM and multiple immunohistochemical markers (Fig. 1) were used to quantify neurons born at the beginning, middle and end of treatments. While a single MEM injection (sMEM) and continuous RUN (cRUN) only transiently increased neurogenesis, extended treatments were capable of increasing neurogenesis at later timepoints. Neurogenic efficacy depended on sex and treatment: in females, 2 months of interval RUN (iRUN) increased DCX^+^ cells; in males, DCX^+^ cells we elevated after 1 month of iRUN followed by 1 month of multiple MEM injections (mMEM). However, thymidine analog labeling revealed that all extended treatments were relatively ineffective at increasing numbers of neurons born in the earlier phases of treatment.

**Figure 1:**
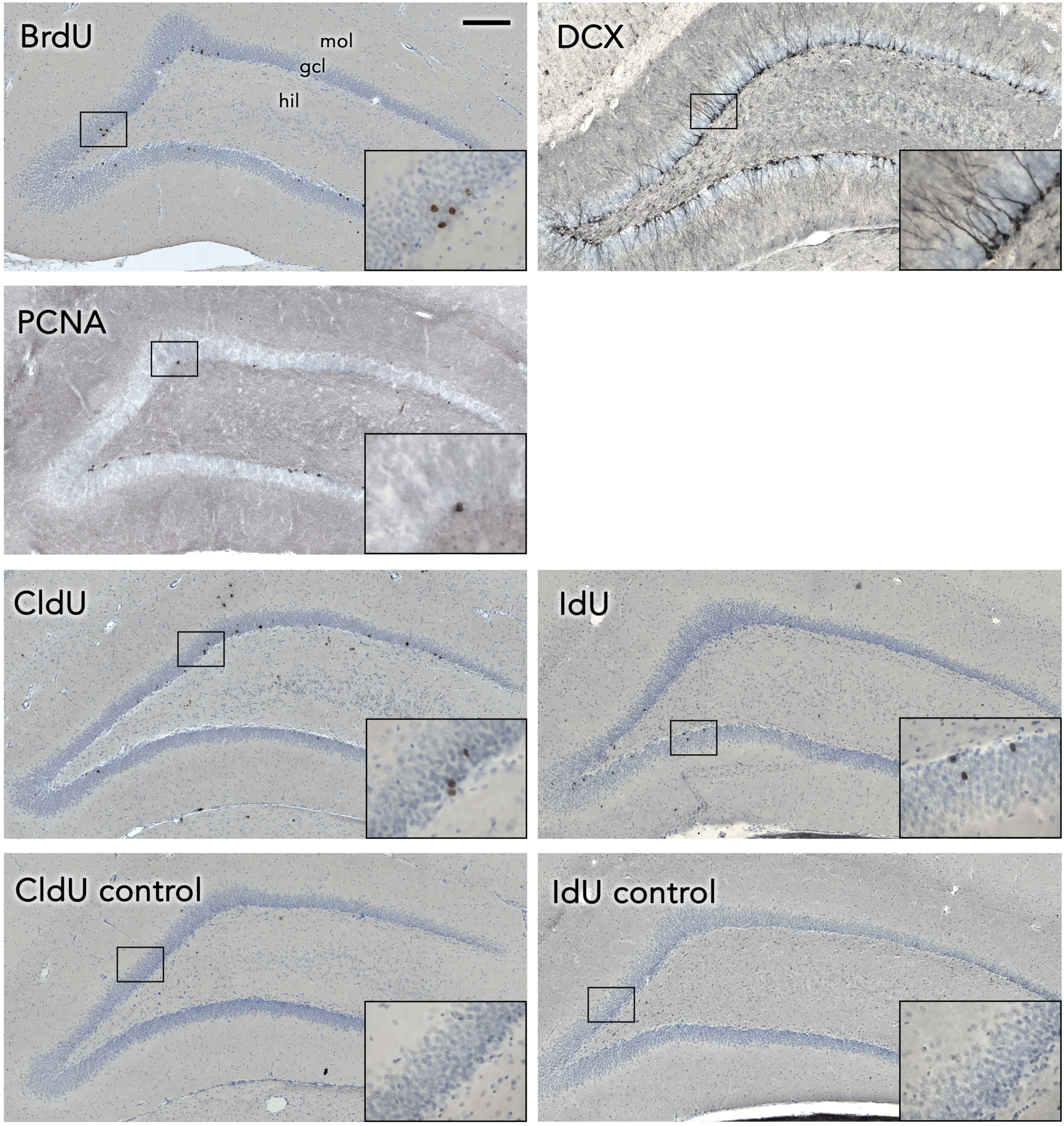
Immunohistochemistry for neurogenesis markers used in this study. Thymidine analogs (BrdU, CldU, IdU) were used to detect neurons born well before the experimental endpoint, DCX was used to detect neurons born in the few weeks preceding the experimental endpoint, and PCNA was used to detect cells that were proliferating at the experimental endpoint. Since CldU and IdU were used in the same animals, and they have the potential to cross react, controls tissue was stained to ensure antibody-antigen specificity (“CldU control” was injected with only IdU and stained for CldU, ie with rat anti-BrdU antibody; “IdU control” was injected with only CldU and stained for IdU, ie with mouse anti-BrdU antibody). Scale bar = 200 μm.

## METHODS

### Animals and Treatments

All procedures were approved by the Animal Care Committee at the University of British Columbia and conducted in accordance with the Canadian Council on Animal Care guidelines regarding humane and ethical treatment of animals. Experimental Long-Evans rats were generated in the Department of Psychology‘s animal facility with a 12-hour light/dark schedule and lights on at 6:00am. Breeding occurred in large polyurethane cages (47cm × 37cm × 21cm) containing a polycarbonate tube, aspen chip bedding and ad libitum rat chow and water. Breeders (both male and female) remained with the litters until P21, when offspring were weaned to 2 per cage in smaller polyurethane bins (48cm × 27cm × 20cm) with a single polycarbonate tube, aspen chip bedding and ad libitum rat chow and tap water. In Experiment 3, animals housed on a reverse light dark cycle (beginning at weaning) in order to study running effects during the active cycle, when rats are most active and neurogenic effects are greatest (Holmes et al., 2004; van der Borght et al., 2006). Running wheel cages consisted of 24” x 18“ x 15” plastic tub containing aspen chip bedding, ad libitum rat chow, water and a 12” running wheel (Wodent Wheel, Exotic Nutrition). Running distance was measured by attaching a neodymium magnet to the running wheel, which allowed each revolution to be detected by a bicycle odometer positioned on the outside of the cage.

This study is comprised of 3 experiments examining the effects of different methods of increasing adult neurogenesis in both males and females. Treatments started at 2 months of age for all experiments.

In **Experiment 1** animals were either given continuous running wheel access (cRUN group) or a single memantine treatment (sMEM group; see timeline in Fig. 2A) and compared to their respective controls. cRUN animals were pair housed and given free access to running wheels in their home cages. cRUN animals were compared to sedentary controls that were pair housed without access to a running wheel. Seven days after cRUN treatment began, animals were given a single BrdU injection (200 mg/kg, I.P.; Sigma, cat #B500205). sMEM animals were given a single memantine injection (35 mg/kg, I.P; Toronto Research Chemicals, cat #M218000100) at 2 months of age followed 3 days later by a single BrdU injection (200 mg/kg, I.P.). sMEM animals were compared to vehicle-injected controls. All animals were perfused with 4 % paraformaldehyde 4 weeks after BrdU injection and brains were extracted and post-fixed for an additional 48 hours.

**Figure 2:**
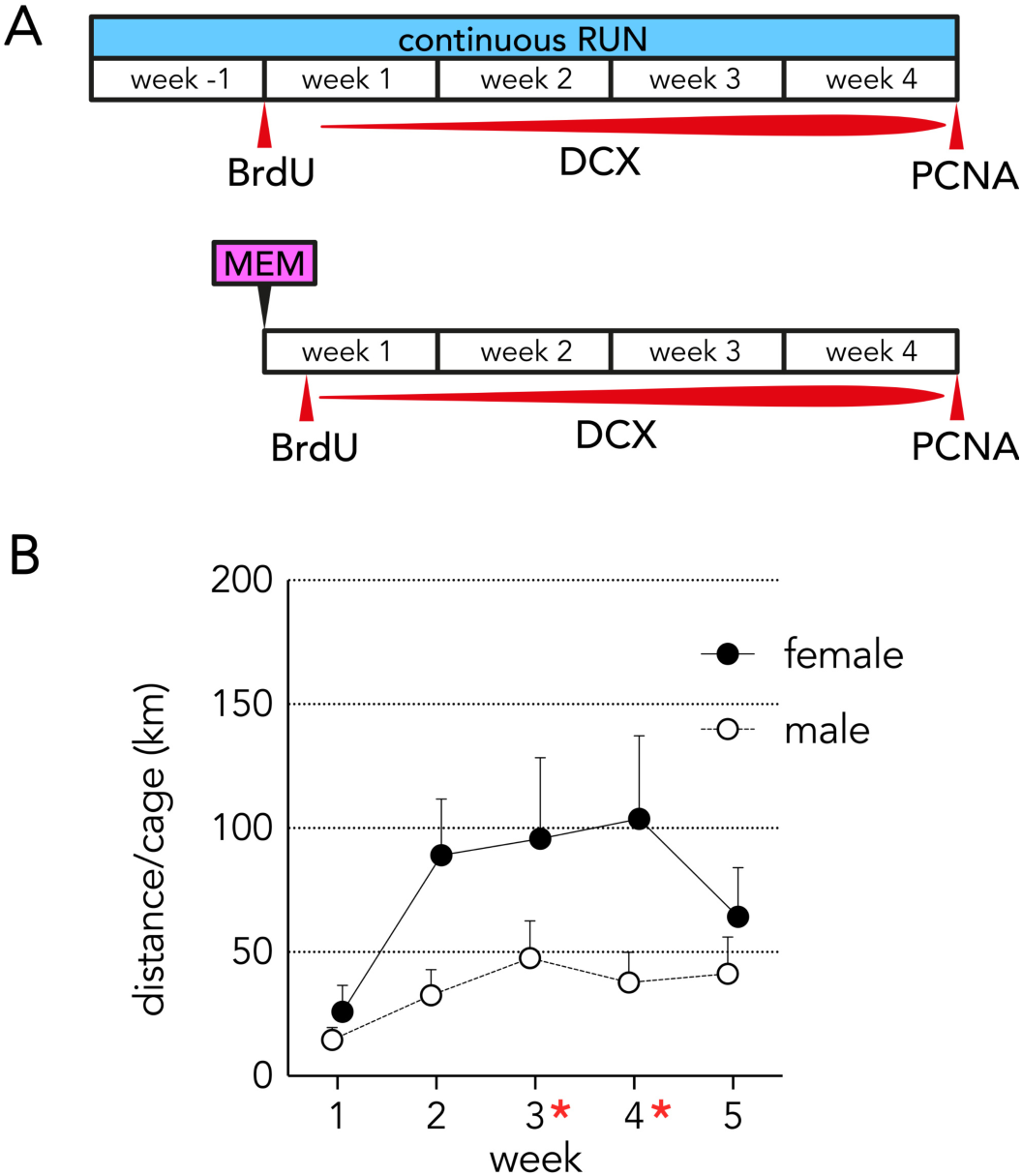
Experiment 1: Short-term treatment with cRUN and sMEM. A) Experimental timeline: cRUN rats were given continuous access to running wheels and sMEM rats were given a single injection of memantine. A single BrdU injection labelled a discrete population of cells born in the early phase of treatment, PCNA immunostaining was used to visualize cells that were actively proliferating at the end of the experiment when the animals were killed, and DCX immunostaining was used to visualize neurons born in the (primarily 2) weeks prior to death. B) Average running distance per cage (i.e. pair of cRUN animals) increased over time (bars indicate standard error; *P<0.05 vs 1w).

In **Experiment 2** we assessed the long-term effects of continuously housing male rats with access to a running wheel. This was the only experiment that did not include female rats. These animals were injected with BrdU (50 mg/kg, I.P.) on postnatal day 6 as a part of a separate study. At 2 months of age rats were pair housed with constant access to running wheels for 4 months, and were perfused with 4% paraformaldehyde at 6 months of age.

In **Experiment 3** groups were given 2 × 4-week treatment blocks according to 5 possible combinations: CON, iRUN/iRUN, mMEM/mMEM, iRUN/mMEM, mMEM/iRUN (see timeline in Fig. 6A). CON rats remained in their home cages throughout both treatment blocks, and were handled by an experimenter on MEM injection days. A block of mMEM treatment consisted of 4 weekly MEM injections (35 mg/kg each). A block of iRUN treatment consisted of rats being placed individually in running wheel cages for 4 hours on weekdays. The iRUN was counterbalanced, so that rats would run for the first four hours of the dark phase on one day and the middle four hours of the dark phase on the next day. On weekends (3 nights/week), iRUN rats were pair housed in the running wheel cage, with free access to the wheels. Thymidine analogues (CldU and IdU, see below Fig 6A) were used to label neurons born at the beginning of the first and second block of treatments, respectively. For mMEM blocks, the thymidine analog was injected 3 days after the first MEM injection. For the iRUN blocks, the thymidine analogue was injected 5 days after first 4-hour block of running. For the first treatment block, whether iRUN or mMEM, rats were injected with CldU (42.5 mg/kg, IP; Toronto Research Chemicals, cat #2105478) and for the second treatment block, whether iRUN or mMEM, rats were injected with IdU (57.5 mg/kg, IP; Toronto Research Chemicals, cat #2100357)(Vega & Peterson, 2005). Immediately following the second block of treatment animals were perfused with 4% paraformaldehyde and brains were extracted and post-fixed for an additional 48 hours.

### Tissue processing and immunohistochemistry

Brains were immersed in 10% glycerol solution for 1 day, 20% glycerol solution for 2 days and then sectioned coronally at 40 μm on a freezing microtome. Sections were stored in cryoprotectant at -20°C until immunohistochemical processing. To detect BrdU^+^, CldU^+^ or IdU^+^ cells in a 1 in 12 series of sections throughout the entire dentate gyrus were mounted onto slides, heated to 90°C in citric acid (0.1M, pH 6.0), permeabilized with trypsin, incubated in 2N HCl for 30 min to denature DNA, and incubated overnight at 4°C with mouse anti-BrdU antibody (to detect BrdU and IdU; BD Biosciences; 347580, USA) or rat anti-BrdU antibody (to detect CldU; BioRad, OBT0030G, USA). Anti-BrdU antibodies were diluted 1:200 in 10% triton-x and 3% horse serum. Sections were washed and incubated in biotinylated goat anti-mouse (Sigma, cat #B0529) or biotinylated donkey anti-rat (Jackson, cat #712065153) secondary antibody for 1 hour (1:250), tissues was blocked in 0.3% H_2_O_2_ for 30 min, and cells were then visualized with an avidin-biotin-horseradish peroxidase kit (Vector Laboratories, cat #OK-6100) and cobalt-enhanced DAB (Sigma Fast Tablets, cat #DO426). Sections were then rinsed in PBS, dehydrated, cleared with citrisolv (Fisher, cat #22143975) and coverslipped with Permount (Fisher, cat #SP15500). To detect PCNA^+^ cells a 1 in 12 series of sections throughout the entire dentate gyrus were mounted onto slides, heated to 90°C in citric acid (0.1 M, pH 6.0), permeabilized with trypsin, incubated in 2N HCl for 30 min, incubated overnight at 4 °C with mouse anti-PCNA antibody (1:200, Santa Cruz Biotechnology, cat #sc-56) and then processed as per thymidine analogs. Two sections containing the dorsal hippocampus were stained for the immature neuronal marker doublecortin (DCX) to detect immature neurons. In Experiment 1, staining was performed on free-floating sections with fluorescent detection. Sections were treated with PBS with 10% triton-x, 3% horse serum for 30 minutes, incubated at 4°C for 3 days in PBS with 10% triton-x, 3% horse serum and goat anti-DCX (1:250; Santa Cruz Biotechnology, cat #sc-8066). Visualization was performed with Alexa 555-conjugated donkey secondary antibody (Invitrogen/Thermofisher, cat #A21432) diluted 1:250 in PBS for 60 minutes at room temperature. Sections were counterstained with DAPI (Life Technologies, cat #D1306), mounted onto slides and coverslipped with PVA-DABCO. In experiment two, sections were mounted on slides, heated to 90°C in citric acid (0.1 M, pH 6.0), sections were washed, permeabilized in PBS with 10% triton-x for 30 min and incubated for three days at 4 °C with goat anti-DCX (1:250 in 10% triton-x and 3% horse serum). Sections were washed and incubated in biotinylated donkey anti-goat secondary antibody for 1 hour (1:250, Jackson, cat #705065147) and processed for peroxidase-DAB as above.

### Microscopy and sampling

Quantification of all DAB-stained cells was performed using a brightfield Olympus CX41 microscope and a 40x objective. For BrdU, IdU, CldU and PCNA, a 1 in 12 series of sections spanning the entire dentate gyrus was examined and all cells that were located within the granule cell layer or its 20 μm hilar border (the subgranular zone) were counted and multiplied by 12 to estimate the total number of cells per dentate gyrus (bilaterally). DAB-stained DCX^+^ cells were quantified across the entire granule cell layer and subgranular zone (∼20 μm wide) from 2 dorsal sections (4 hemispheres). The granule cell layer volume was calculated by multiplying the section thickness (40 μm) by the 2D area (measured from 2x images with ImageJ (NIH)), which was then used to calculate DCX^+^ cell densities.

### Statistical Analyses

Unpaired t-tests and two-way (sex x treatment) ANOVAs were used to detect effects of treatment and sex on measures of neurogenesis. Where significant interactions were observed, Sidak’s (Experiment 1) or Dunnett’s tests (Experiment 3) were used to compare treatment groups to controls. In cases where distributions failed tests of normality and homogeneity of variance, analyses were run on log transformed data or non-parametric tests were used as described below. In all cases statistical significance was set at p = 0.05.

## RESULTS

### Experiment 1 – Short term continuous RUN and a single MEM injection

#### Short-term continuous running behavior

Rats housed with running wheels ran progressively more over time (Fig. 2B; effect of time: F_4,24_= 3.8, P= 0.015). Since rats were pair housed, we could only determine the distance run per cage rather than per individual rat. This limitation precluded a thorough examination of sex differences but, nonetheless, female pairs tended to run more than males (effect of sex F_6,24_= 4.6, P=0.09).

#### Five weeks of continuous running transiently increases neurogenesis

Neurogenesis was measured with 3 immunohistochemical markers that label complementary populations of cells born at different phases of the cRUN treatment. BrdU^+^ cells, born after the first week of running, were increased in both males and females (log transformed data: effect of RUN F_1,26_=26.8, P<0.0001; Fig. 3A). There tended to be more BrdU^+^ cells in males, and the cRUN-induced increase in BrdU^+^ cells was twice as large in females as in males (84% vs 42%), but sex differences were not statistically different (effect of sex F_1,26_=3.4, P=0.08; interaction F_1,26_=2.4, P=0.13). DCX^+^ cells were quantified to examine cumulative cRUN effects on neurons mainly born during the last 2 weeks of running (Brown et al., 2003; Rao & Shetty, 2004; Snyder et al., 2009). Here, we also found that DCX was higher in runners compared to sedentary rats (effect of cRUN F_1,28_=4.6, P=0.04; Figure 3B). However, the neurogenic effect of running was weaker than for BrdU^+^ cells, with only a 14% increase in males and 12% increase in females. PCNA immunostaining, to identify cells that were proliferating at the end of the 5 weeks of running, revealed modest increases in runners compared to sedentary controls (effect of running, F_1,27_=3.8, P=0.06; 38% increase in males, 14% increase in females; Fig. 3C). This experiment demonstrates that running initially increases neurogenesis but any effects remaining after 5 weeks are marginal.

**Figure 3:**
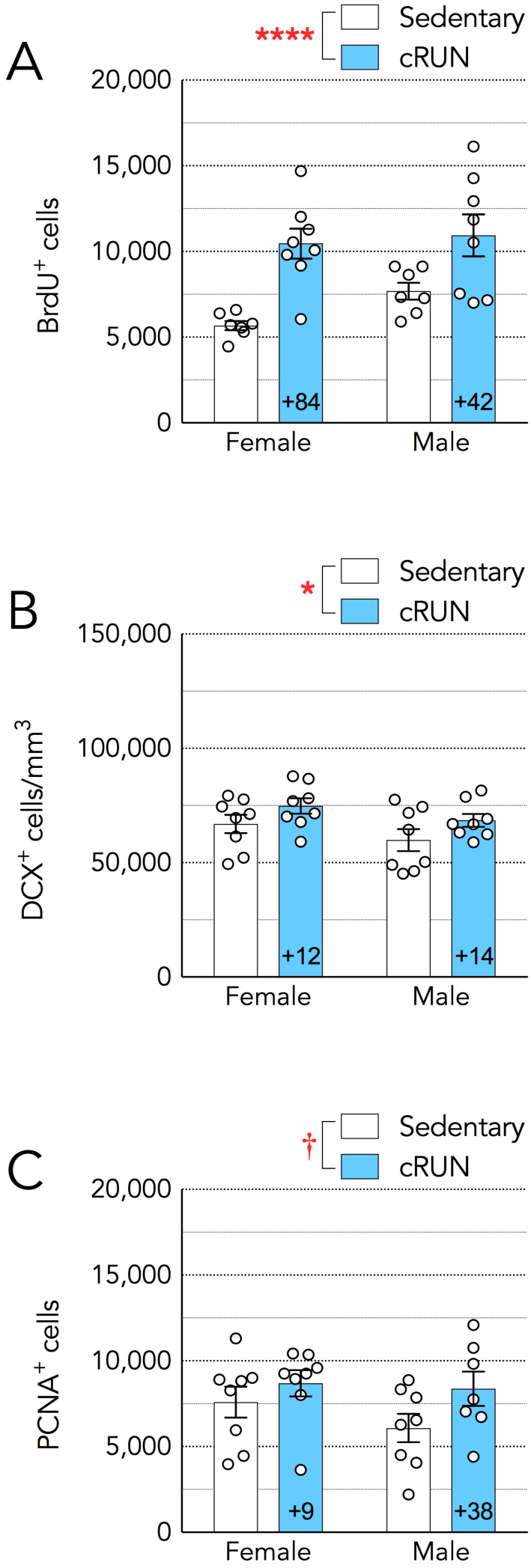
Experiment 1: Transient effects of running on neurogenesis. A) Five weeks of running increased neurogenesis as measured by BrdU labelling of neurons born after the first week of running. B) DCX^+^ cells, reflecting neurons born in the last ∼3 weeks, were increased in runners by < 15%. C) PCNA^+^ cells, proliferating at the end of 5 weeks of cRUN, were not significantly increased in runners compared to sedentary controls. ****P<0.0001, *P<0.05, 0.05 < ^†^P <0.1. Graphs indicate group means ± standard error. Numbers embedded in cRUN bars indicate running effect size (%) relative to sedentary controls.

#### A single MEM injection transiently increases neurogenesis

MEM injection increased the number of BrdU^+^ cells that were born 3 days later and survived for an additional 4 weeks (treatment effect F_1,18_=14.5, P=0.0013; Fig. 4A). The relative increase in BrdU^+^ cells was identical in males and females (60% over controls). There were no differences in numbers of DCX^+^ or PCNA^+^ cells due to sMEM or sex (Fig. 3B,C). Thus, as with running, the neurogenic effects of sMEM were transient.

**Figure 4:**
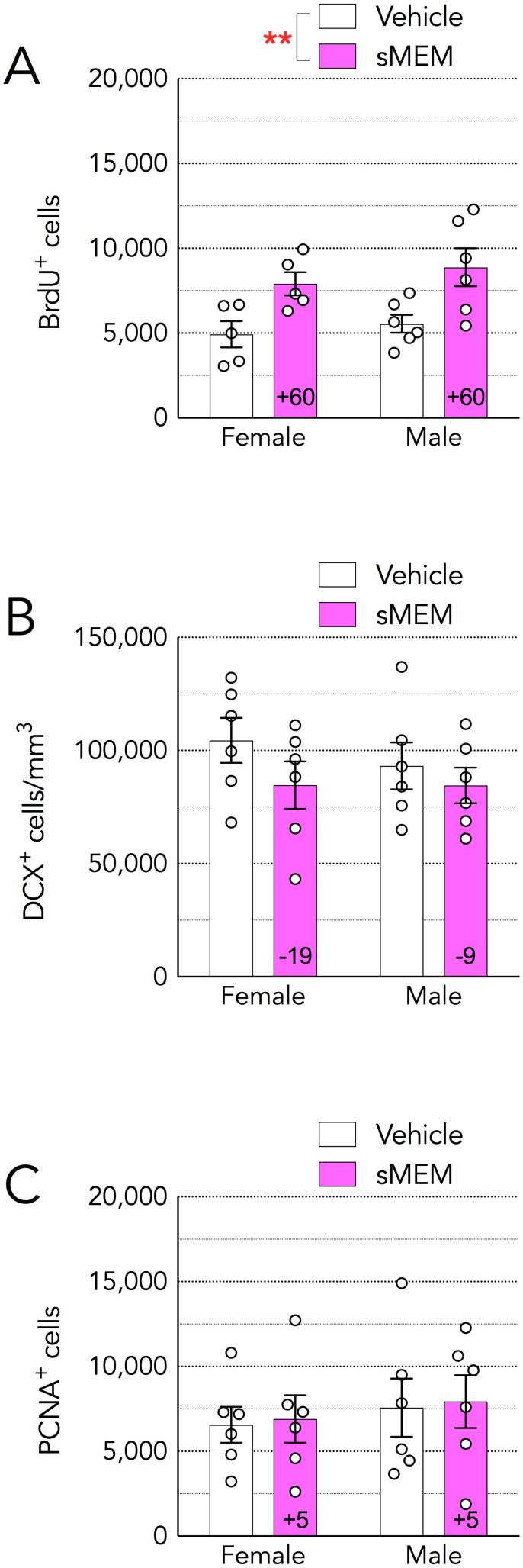
Experiment 1: Transient effects of a single memantine injection on neurogenesis. A) sMEM increased the number of BrdU^+^ cells born 3 days later and surviving for ∼ 4 more weeks. B) Four weeks after sMEM, numbers of immature DCX^+^ cells were not different from vehicle-injected controls. C) Four weeks after sMEM, numbers of proliferating PCNA^+^ cells were equivalent to vehicle controls. **P<0.01. Graphs indicate group means ± standard error. Numbers embedded in sMEM bars indicate effect size (%) relative to vehicle-injected controls.

### Experiment 2 – Long-term continuous RUN

#### Long-term continuous running behavior

Since rats were pair housed running data reflects activity per cage and not per animal. Tracking for one cage was incomplete and therefore not included. For the remaining 3 cages, running increased over the first 3 weeks and then subsided to stable levels for the remaining weeks (Fig. 5A; repeated measures ANOVA, effect of time F_2,56_=2.4, P=0.011; week 3 vs weeks 1,14-19 all P<0.05).

**Figure 5:**
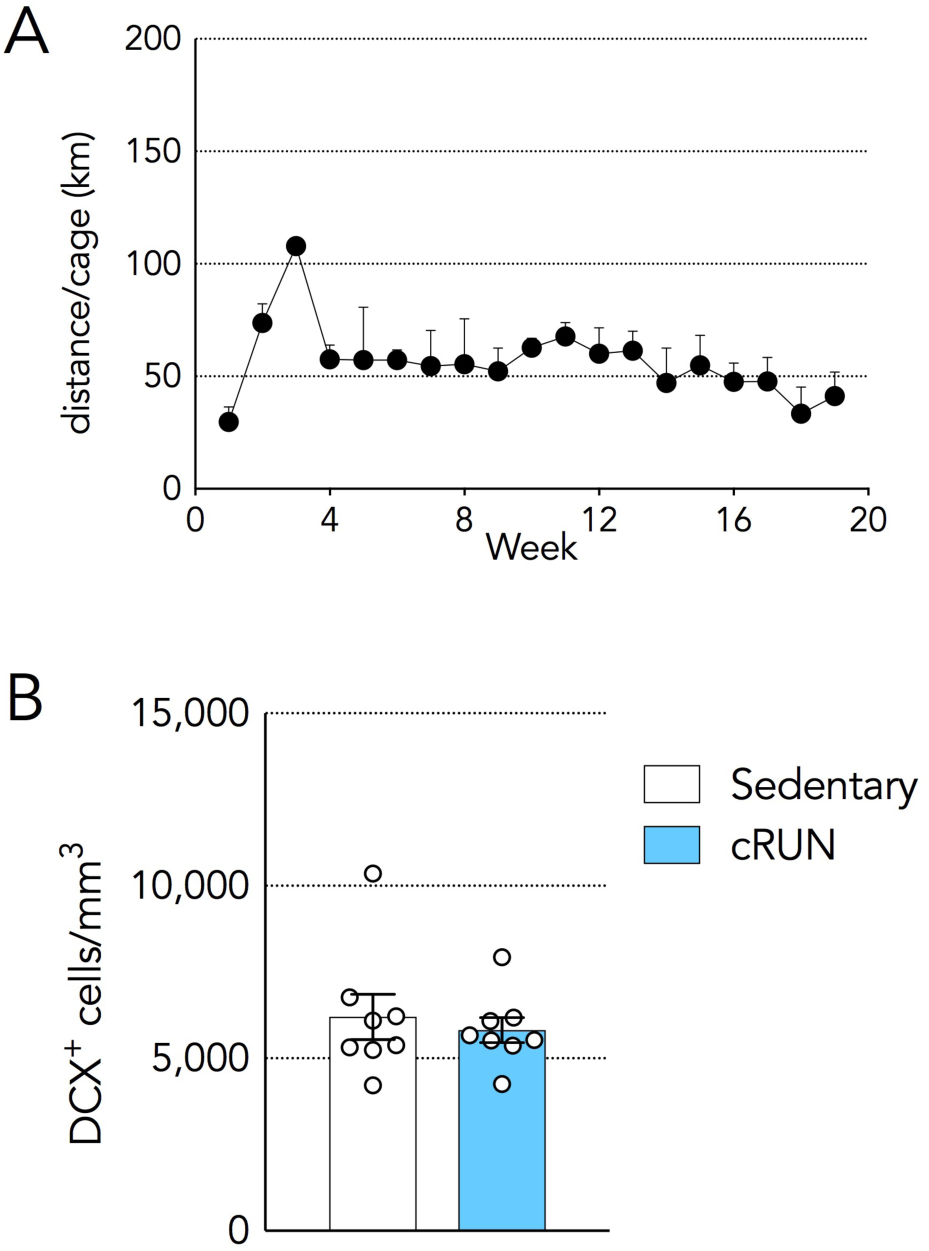
Experiment 2: Long-term continuous running. A) Running increased over the first 3 weeks and then decreased to stable levels. B) Numbers of immature DCX^+^ cells were similar in runners and sedentary controls. Symbols and bars indicate mean ± standard error.

#### Effects of long-term continuous running on neurogenesis

To investigate whether the marginal increase in DCX^+^ cells observed after 1 month of cRUN (Experiment 1) persists if cRUN is extended to 4 months, we quantified DCX^+^ cells in cRUN and sedentary rats. We observed no difference between groups, indicating that cRUN treatment does not result in sustained increases in neurogenesis (Fig. 5B; T_14_=0.51, P=0.6).

### Experiment 3 – Long-term interval RUN and multiple MEM injections

#### Long-term interval running behavior

To test whether other paradigms were capable of leading to sustained increases in neurogenesis, rats were subjected to interval RUN (iRUN), multiple MEM injections (mMEM), or combined iRUN + mMEM treatment (Fig. 6A). Rats ran progressively more with each week of iRUN (Fig. 6B). Within-sex ANOVAs revealed that the effect of time was only significant in females (all male groups P> 0.2; female iRUN/iRUN F_7,56_=2.7, P<0.05; weeks 6,7,8 vs week 1, P<0.05; female iRUN/mMEM F_3,39_=8.8, P=0.002; weeks 2,3 vs week 1, P<0.05 and week 4 vs week 1, P<0.01; female mMEM/iRUN F_3,31_=7.2, P=0.007; week 4 vs week 1, P<0.01). An analysis of average weekly running distance across treatment groups and sexes revealed no differences between treatment groups but significantly greater distances run by females (group effect F_2,52_=1.9, P=0.2; sex effect F_1,52_=15, P<0.001; Fig. 6C).

**Figure 6:**
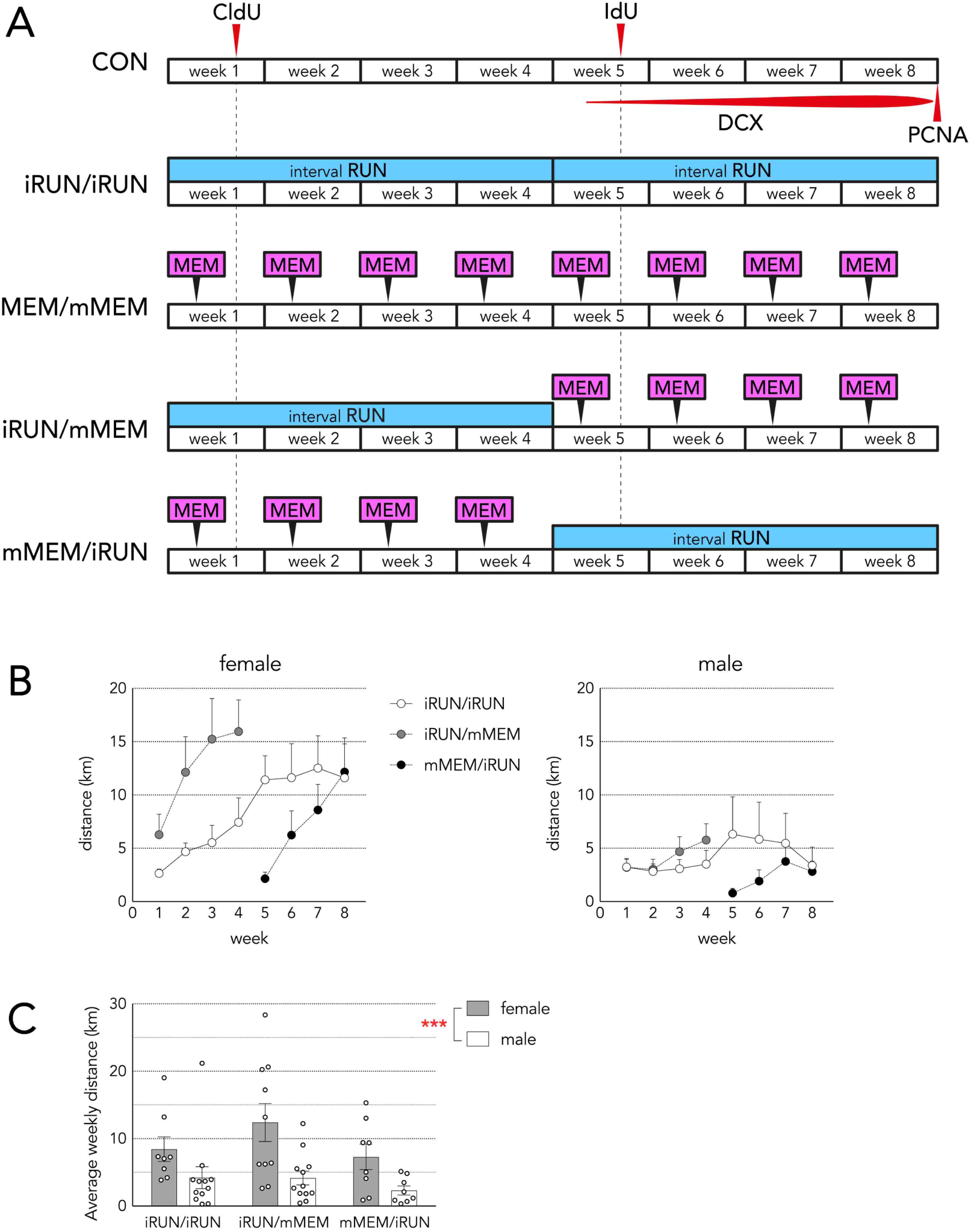
Experiment 3: long-term treatment with iRUN and mMEM. A) The experimental design consists of 2 × 4-week treatment blocks. Four neurogenesis markers were used to identify neurons born at different stages of the 2 treatment blocks: CldU (neurons born at the beginning of block 1), IdU (neurons born at the beginning of block 2, DCX (neurons born during the latter weeks of block 2) and PCNA (neurons born at the end of block 2). CON rats remained in standard cages and were handled on MEM injection days. iRUN/iRUN rats were given 8 weeks of interval running. mMEM/mMEM rats received 8 weekly injections of memantine. iRUN/mMEM rats received 4 weeks of interval running and then 4 weeks of memantine injections. mMEM/iRUN rats received 4 weeks of memantine injections and then 4 weeks of interval running. B) Average weekly running behavior for each group, broken down by sex. Only females ran significantly more with time (see text for details). Bars indicate standard error. C) Average weekly running behavior. Females ran significantly more than males (***P<0.001).

### Effects of long-term mMEM, iRUN and mMEM+iRUN treatments on neurogenesis

To investigate the efficacy of long-term neurogenic manipulations, we used complementary immunohistochemical markers to quantify: neurons born at the beginning of treatment block 1 (CldU), neurons born at the beginning of treatment block 2 (IdU), immature neurons born during the latter portion of treatment block 2 (DCX) and cell proliferation at the end of treatment block 2 (PCNA) (Fig. 7A). To ensure specificity of the CldU and IdU labelling, CldU-only tissue was stained for IdU and IdU-only tissue was stained for CldU (2 animals for each). No thymidine-labelled cells were observed (Fig. 1).

**Figure 7:**
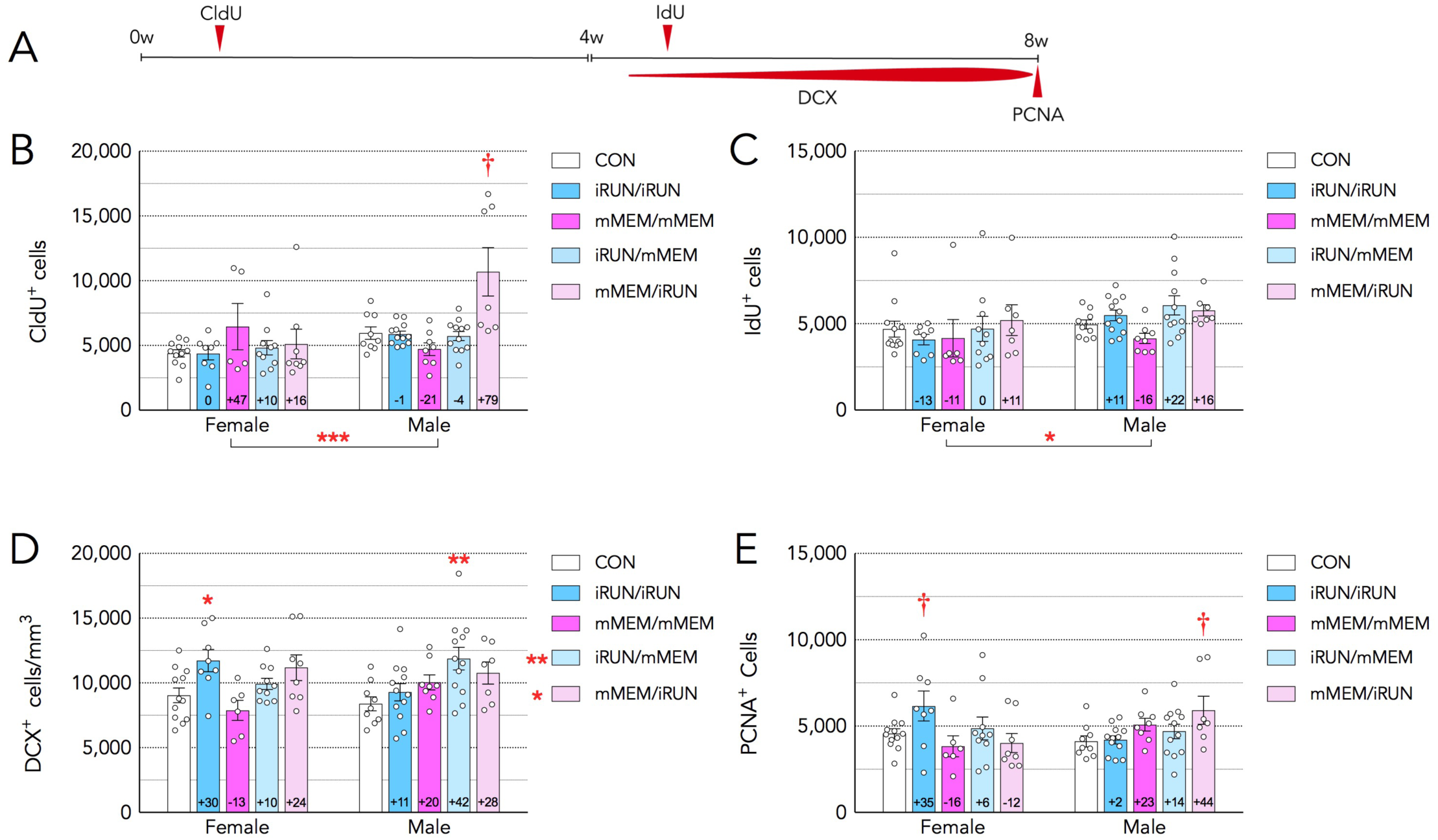
Experiment 3: Effects of long-term iRUN and mMEM treatments on neurogenesis. A) Experimental timeline indicating populations of cells identified by immunohistochemical markers. B) CldU^+^ cells, born at the beginning of block 1, were significantly greater in mMEM/iRUN animals relative to controls. This effect was mainly driven by effects in males. There were also more CldU^+^ cells in males than in females. C) IdU^+^ cells, born at the beginning of block 2, were not different from controls following any of the treatments. There were more IdU^+^ cells in males than in females. D) The density of immature DCX^+^ cells, born during block 2, was greater in the iRUN/mMEM and mMEM/iRUN treatment groups. In males, iRUN/mMEM significantly increased DCX^+^ cell density and, in females, iRUN/iRUN significantly increased DCX^+^ cell density. E) PCNA cell counts revealed that no treatment regimen altered cell proliferation at the end of the experiment. ***P<0.001, **P<0.01, *P<0.05 vs control, 0.05<^†^P<0.1 Dunn’s test vs control). Bars indicate group means ± standard error.

Numbers of CldU^+^ cells, born at the beginning of block 1, were 30% greater in males than females (log transformed data: sex effect F_1,80_=15, P=0.0002; Fig. 7B). There was no main effect of treatment (F_4,80_=2.3; P=0.07) but a sex x treatment interaction (F_4,80_=3.6; P=0.009) where mMEM/iRUN increased CldU^+^ cells relative to controls in males only (79% increase in males, P=0.01; 16% increase in females). However, the individual data points clearly reveal that mMEM/iRUN effects on CldU^+^ cells were variable: 3 of 7 rats had CldU^+^ cell numbers that were approximately twice control levels, the other 4 rats were within the range of controls, and re-analysis with a non-parametric test revealed no difference from controls (Dunn’s test: mMEM/iRUN vs control, P=0.09). Non-parametric analysis of overall CldU^+^ cells in males vs females confirmed greater numbers of cells born in males during the initial phase of treatment (Mann Whitney test, P<0.0001).

To examine cells born at the beginning of the 2^nd^ treatment block, we quantified IdU^+^ cells (Fig. 7C). The total number of IdU^+^ cells was (17%) greater in males than in females (effect of sex F_1,81_=4.1, P=0.046). However, there were no effects of treatment on IdU^+^ cell numbers (effect of treatment F_4,81_=1.6, P=0.17; treatment x sex interaction F_4,81_=0.7, P=0.6). Since these data were not normally distributed, even after transformation, we additionally examined the overall sex difference with non-parametric tests and confirmed that males had more IdU^+^ cells than females (Mann Whitney test, P=0.0003).

To examine a broad-aged population of cells born in the 2^nd^ block of the extended treatments, we quantified immature DCX^+^ neurons (Fig. 7D). Here, we found a significant effect of treatment, where only the combined iRUN+mMEM treatments increased DCX^+^ cell density (across sexes, iRUN/mMEM 26% over controls, P=0.005; mMEM/iRUN 25% over controls, P=0.015). A sex x treatment interaction (F_2,82_= 3.5, P=0.012) revealed distinct patterns in males and females: in females, iRUN/iRUN selectively increased DCX^+^ cell density (30% increase over controls, P=0.03) whereas in males iRUN/mMEM selectively increased DCX^+^ cell density (42% increase, P=0.002).

Finally, to identify whether extended iRUN and mMEM treatments had a sustained impact on neurogenesis, we examined PCNA^+^ cells to determine cellular proliferation levels at the end of the treatment paradigms (Fig. 7E). We found no effect of treatment or sex on proliferating PCNA^+^ cells (treatment effect F_4,82_=0.9, P=0.44; sex effect F_1,82_=0.1, P=0.7). There was a significant sex x treatment interaction (F_4,82_=4.1, P=0.005) and, while differences were not statistically different, the female iRUN/iRUN group had 35% more PCNA^+^ cells than controls (P=0.08) and the male mMEM/iRUN had 44% more PCNA^+^ cells than controls (P=0.07), which is broadly consistent with the elevated DCX^+^ cell densities in these treatment groups.

Numbers embedded in bars indicate effect size (%) relative to controls.

## DISCUSSION

Here we examined the effects of two neurogenic treatments, RUN and MEM, on adult neurogenesis in the dentate gyrus of male and female rats. Our general strategy was to immunostain for thymidine analogs to detect treatment effects on cells born long before the experimental endpoint, DCX to detect effects on neurons born in the weeks prior to endpoint, and PCNA to detect effects on cell proliferation at the very end of each experiment. We have two main findings. First, neurogenic effects were temporally limited: cRUN and sMEM increased neurogenesis only at early timepoints and iRUN and mMEM increased neurogenesis only at the later timepoints. Second, there were sex differences in the neurogenic treatment efficacy: iRUN increased neurogenesis in females and iRUN/mMEM increased neurogenesis in males. Neurogenesis was also greater in males than in females, as measured by thymidine analogs. Collectively, these results identify temporal and sex-dependent constraints on the ability to produce new dentate gyrus neurons in adulthood.

### Transient neurogenic effects of sMEM and cRUN

In Experiment 1 we confirmed previous reports that both cRUN and sMEM can have potent neurogenic effects, increasing numbers of surviving BrdU^+^ cells by 40-80%. BrdU was injected 1 week after cRUN onset, to capture combined neurogenic effects on proliferation and survival (Snyder et al., 2009). In the case of sMEM, BrdU was injected 3 days after MEM injection, to capture the delayed effects of MEM on proliferation (Maekawa et al., 2009). Running showed some indication of lasting effects as both sexes had elevations in DCX^+^ and PCNA^+^ cells at the end of 5 weeks. However, these effects were modest and, in the case of PCNA, not statistically different from controls. Moreover, in Experiment 2 we found no trend for increased neurogenesis following 4 months of cRUN. While RUN can increase neurogenesis beyond 30 days (Patten et al., 2013), many studies have reported decreased effects with time (Clark et al., 2010; Kronenberg et al., 2006; Naylor et al., 2005; Snyder et al., 2009). sMEM effects were also transient, increasing BrdU^+^ cell numbers but without effects on DCX^+^ cells or PCNA^+^ cells that were born days and weeks later. This experiment provides new data on the transient nature of MEM effects, since previous studies have only examined neurogenic effects at short post-MEM intervals (Ishikawa et al., 2014; Maekawa et al., 2009; Namba et al., 2009; Sun et al., 2015).

### Sex differences in the efficacy of extended neurogenic treatments

Given the transient effects of cRUN and sMEM, in Experiment 3 we sought to determine whether iRUN, mMEM, or combined iRUN+mMEM treatments were capable of producing sustained increases in neurogenesis. As reflected by DCX^+^ cell densities, we indeed observed elevated neurogenesis after 2 months of treatments. The only treatments that increased DCX^+^ cells when males and females were pooled (i.e. main effect) were the combined iRUN+mMEM groups. Whereas RUN increases neurogenesis by increasing the division of transit amplifying cells (Kronenberg et al., 2003) and promoting neuronal survival (Snyder et al., 2009), MEM appears to increase neurogenesis by promoting symmetric division, and therefore expansion, of radial glial stem cells (Namba et al., 2009). Thus, by targeting distinct and complementary mechanisms, combined RUN+MEM treatment may be more effective than either treatment in isolation. This may be particularly true for males, as analysis of the sex x treatment interaction revealed that they were more responsive to the neurogenic effects of combined RUN+MEM treatments. Notably, males receiving RUN+MEM treatment also showed a trend for increased proliferation, with 44% more PCNA^+^ cells after iRUN/mMEM.

In contrast, only females responded to the iRUN/iRUN treatment with elevated DCX^+^ cell densities, and a trend for greater proliferation (35% increase). The greater efficacy of RUN in females fits with the general observation that female rodents are more active than males (Lightfoot, 2008) and evidence that female running distance is correlated with the addition of new cells (Rhodes et al., 2003). Notably, in females, 2 months of iRUN increased DCX^+^ cell density more than 1 month of cRUN. This is despite the fact that they ran less in the iRUN condition than in cRUN condition (∼2 vs ∼5 km/rat/day, respectively). This aligns with a number of other studies that have found that restricted, or less intense, running has greater neurogenic effects than continuous, or more intense, running (Grégoire et al., 2014; Inoue et al., 2015; Naylor et al., 2005; Nguemeni et al., 2018; So et al., 2017). The lack of effect of iRUN on males could reflect different neurogenic regulatory mechanisms or it could simply be because they did not run sufficient distances (∼0.5 km per daily 4-hour session). While forced running paradigms could be used to equalize exercise across sexes, these manipulations introduce new confounds since they differentially impact neurogenesis and anxiety levels (Leasure & Jones, 2008). As noted by others, these types of findings warrant additional research into the neurobiological effects of exercise in males vs. females and they indicate that, as a therapy, exercise likely needs to be tailored to individual needs (Barha, Galea, Nagamatsu, Erickson, & Liu-Ambrose, 2017).

### Limited neurogenic effects of RUN and MEM

While extended treatments were capable of increasing neurogenesis at the end of the 2 months (as seen primarily with DCX), they were less effective at increasing neurogenesis at the earlier stages of treatment (as detected by CldU and IdU). In the case of RUN, our female data are therefore consistent with recent work showing that unrestricted exercise promotes neurogenesis at early but not late timepoints, and interval/restricted exercise promotes neurogenesis at late but not early timepoints (Nguemeni et al., 2018).

Surprisingly, while sMEM increased neurogenesis in both females and males, mMEM was much less effective. First, MEM increased BrdU^+^ neurons that were born 3 days after the first MEM injection in Experiment 1 but it did not reliably increase CldU^+^ or IdU^+^ neurons that were born 3 days after the first MEM injection in Experiment 3. This discrepancy suggests that a single MEM injection increases the birth of new neurons but subsequent MEM injections reduce the survival of those neurons. Second, more cells were born 3 days after a single injection of MEM in Experiment 1, but not 3 days after the 4^th^ or 8^th^ injection in Experiment 3. This indicates that the proliferative effects of MEM dissipate with additional treatments. Nonetheless, males seemed to show relatively greater benefit from mMEM treatment: iRUN/mMEM males had increased DCX^+^ cell densities, and mMEM/iRUN males showed a trend for more CldU^+^ cells.

The limited efficacy of MEM could result from homeostatic mechanisms whereby an injection of MEM increases neurogenesis which in turn reduces both the survival of cells born prior to the injection and the proliferation of cells in the weeks after the injection. Alternatively, sMEM and mMEM could impact neurogenesis through a multitude of other mechanisms. Indeed, positive effects of MEM on hippocampal BDNF levels are observed after acute but not chronic treatment (Réus et al., 2010). Additionally, while MEM is therapeutically attractive given its positive effects on hippocampal-dependent learning (Ishikawa et al., 2014; Zajaczkowski, Quack, & Danysz, 1996; Zoladz et al., 2006) and neuroprotective functions (Creeley, Wozniak, Labruyere, Taylor, & Olney, 2006), it alters the migration of adult-born neurons (Namba et al., 2011), and may have negative effects under certain treatment regimens. Negative effects may be more pronounced in females as reported for other NMDAR antagonists (Fix et al., 1995). While MEM has been reported to be relatively innocuous compared to MK-801, MEM concentrations in serum and brain are twice as high in females as compared to males after intraperitoneal injection (Zajaczkowski et al., 2000). These sex differences broadly fit with our observation that mMEM treatment retained some neurogenic effects in males but not females.

### Implications for neurogenic therapies

Our findings have a number of implications for disorders that impact the structural integrity of the hippocampus. Alzheimer’s disease, schizophrenia and depression differentially impact males and females (Aleman et al., 2003; Gao et al., 1998; Seedat et al., 2009) and all are associated with hippocampal volume loss (Harrison, 2004; Jack et al., 2000; McKinnon et al., 2009). Our results clearly show that there are sex differences in the efficacy of treatments that increase neurogenesis and could potentially offset structural deficits. While both RUN and MEM were capable of increasing neurogenesis in both sexes, iRUN was more effective in females and combined RUN+MEM treatments were more effective in males. However, both short and long treatments were only partially effective, with short treatments increasing neurogenesis only at early timepoints and longer treatments increasing neurogenesis only at later timepoints. Additional work is needed to identify whether extended treatment paradigms can increase neurogenesis beyond 2 months. If not, this may suggest that there are constraints on the extent to which neurogenesis can be increased, which could be due to a limited capacity of the stem cell pool (Encinas et al., 2011). Treatments that promote stem cell renewal may therefore be particularly fruitful for ensuring the long-term production of new neurons. Finally, given the plastic potential of adult-born neurons, it is equally important to consider treatments that exploit their physiological functions, which could exert powerful effects on circuits and behavior, even in the absence of large changes in neuronal numbers.

## ACKNOWLEDGEMENTS

This work was supported by an NSERC Discovery Grant (JSS), an NSERC postgraduate scholarship (SPC) and Killam Doctoral Scholarship (SPC).

